# Spatial transcriptomic analysis of progressing oral epithelial dysplasia reveals unique differentially expressed genes and microenvironmental changes

**DOI:** 10.64898/2026.01.07.697832

**Authors:** Vincent Lavoie, Will Jeong, James S. Jeon, Juan F. Andrade, Aiman Ali, Graziella Rigueira Molska, Najmeh Esfandiari, Hui Ling Yeo, Igor Jurisica, Deepika Chugh, Iona Leong, Justin Bubola, Grace Bradley, Marco Magalhaes

## Abstract

Oral squamous cell carcinoma (OSCC) often arises from oral epithelial dysplasia (OED); however, the gene expression changes during OED progression and its microenvironment are not fully understood. This study used spatial transcriptomics to identify differentially expressed genes and microenvironmental alterations associated with OED’s malignant transformation of OED. A ten-year retrospective analysis of paired OSCC and prior OED samples was conducted at the University of Toronto Oral Pathology Laboratory. A total of 24 paired progressing OED cases and 23 matched non-progressing OED cases were examined using spatial transcriptomics in PanCK+ (dysplastic epithelium or OSCC) and PanCK- (stroma) regions. The analysis included differential gene expression, pathway analysis and spatial deconvolution. Three genes (STOM, KIF26A, and CDKN2A) showed increased expression in the epithelial component of progressing OED compared with non-progressing OED, whereas 41 genes were differentially expressed in OSCC versus the precursor samples. Ubiquitination-related pathways were enriched during OED progression. Functional validation identified TNFRSF12A (Fn14) as a potential regulator of OED progression to OSCC. The OSCC microenvironment displayed increased numbers of fibroblasts, neutrophils, monocytes, and mast cells compared with that of the precursor samples. Our findings suggest that spatial profiling of OED can help identify unique gene signatures and microenvironmental changes that occur before the malignant transformation.

## Introduction

In 2020, lip and oral cavity cancers ranked as the 16th most common cancer type, accounting for 2.0% of all new cancer cases worldwide (Sung et al., 2021). The lifetime risk is approximately 0.48%, with men being more frequently affected than women (Sung et al., 2021). Among these, oral squamous cell carcinoma (OSCC) accounts for 77-90% of oral cavity cancers (Abadeh et al., 2019). In 2020, an estimated 5400 Canadians developed oral cancer, and 1,500 died from the disease (Brenner et al., 2020). The five-year survival rate is approximately 64% (Brenner et al., 2020). Unfortunately, OSCC is associated with significant morbidity and mortality, and early detection is crucial for improving patient outcomes.

The development of OSCC may be preceded by an oral potentially malignant disorder (OPMD), such as oral leukoplakia (OL) and oral erythroleukoplakia (OEL). The WHO recognizes a group of 11 clinically defined entities and additional familial cancer syndromes, some of which may exhibit oral epithelial dysplasia (OED) upon histological examination. The worldwide prevalence of OPMDs is estimated to be 4.47%, with geographic regions in Asia, South America, and the Caribbean being the most affected (Mello et al. 2018). The reported risk of malignant transformation (MT) of OPMDs varies widely, ranging from 0.13-40.8% across previous studies (Mello et al., 2018; Napier & Speight, 2008). Therefore, multiple studies have investigated the potential clinical, histopathological, and molecular determinants of MT in OSCC.

Recent technological advances have enabled spatially resolved high-throughput sequencing of cancer and precancerous specimens (William et al., 2019). To our knowledge, no study has reported transcriptomic data from paired OED and OSCC samples, based on distinct epithelial, stromal, and inflammatory infiltrate tissue components. Therefore, we characterized the protein-coding transcriptome of progressing and non-progressing OED. Such studies may help establish the molecular, biological, and microenvironmental changes associated with MT and identify biomarkers that could be used for prognosis and clinical management.

## Materials and Methods

### Case selection and pathology review

This retrospective study was approved by the Research Ethics Board of the University of Toronto (protocol #40829). The cases were retrieved from the archives of the Toronto Oral Pathology Service of the Faculty of Dentistry, University of Toronto. A database search identified sequential biopsies of hyperkeratosis or OED, with or without malignant transformation into OSCC. The inclusion criteria were sequential samples from the same intraoral anatomical site and a sufficient sample size (≥ 0.2 cm). The exclusion criteria were labial anatomical site, formalin-fixed paraffin-embedded (FFPE) tissue block age >10 years, specimen fragmentation, and histologic features of HPV-associated OED. Patients were classified as progressors if they developed squamous cell carcinoma ≥ 6 months after their initial biopsy. They were classified as non-progressors if no OED grade change or OSCC development was observed during a period of at least 60 months. A third group of inflamed and non-inflamed non-dysplastic specimens, composed of equal proportions of fibroepithelial polyps and inflammatory fibroepithelial hyperplasia, was included as controls.

Cases from the progressing, non-progressing, and non-dysplastic groups were matched based on demographic and clinicopathological data, including patient sex, age, tobacco consumption history, anatomical site, three-tier WHO oral epithelial dysplasia grade (if applicable), and FFPE tissue block age (Table 1). Post hoc statistical testing for confounding factors included parametric and non-parametric tests using IBM SPSS Statistics version 28 (Chicago, Armonk, NY, USA).

**Table 1:** Clinicopathologic features of progressing and non-progressing groups.

| Clinicopathologic features |  |  |  |  |
| --- | --- | --- | --- | --- |
|  | Variable | Progressing (N=24) | Non-progressing (N=23) | p-value |
| Age (biopsy 1) | Mean (SD) | 60.17 (18.72) | 58.35 (10.69) | 0.686<sup>a</sup> |
|  | Median (min, max) | 63 (15, 86) | **56 (35,79)** |  |
| Age (biopsy 2) | Mean (SD) | 62.39 (18.74) | 61.43 (10.26) | 0.831<sup>a</sup> |
|  | Median (min, max) | 64 (19, 88) | **61 (39, 81)** |  |
| Sex | Male | 15 | 9 | 0.385<sup>b</sup> |
|  | Female | 9 | **12** |  |
| Diagnosis (biopsy 1) | Hyperkeratosis | 2 | 3 | 0.629<sup>c</sup> |
|  | **Low-grade dysplasia** | 7 | **9** |  |
|  | High-grade dysplasia | 15 | 11 |  |
| Anatomical site | **Tongue** | 16 | **16** | **0.607<sup>c</sup>** |
|  | Gingiva | 6 | 3 |  |
|  | **Floor of mouth** | 1 | **3** |  |
|  | Palate | 1 | 1 |  |
| Smoking history | **Current** | 1 | **8** | **0.016<sup>c</sup>** |
|  | Former | 6 | 3 |  |
|  | **Never** | 14 | **12** |  |
|  | Not reported | 3 | 0 |  |
| Block age (biopsy 1) | **Mean (SD)** | 6.50 (1.79) | **6.74 (1.36)** | **0.608<sup>a</sup>** |
|  | Median (min, max) | 4 (2,10) | 4 (5,9) |  |
| Block age (biopsy 2) | **Mean (SD)** | 4.09 (1.50) | **3.91 (1.50)** | **0.697<sup>a</sup>** |
|  | Median (min, max) | 4 (1,6) | 4 (1,7) |  |
| a. Independent samples t-test (2-sided)<br>b. Fischer's Exact Test<br>c. Fischer-Freeman-Halton Exact Test |  |  |  |  |

A blind histopathological review of progressing and non-progressing specimens was performed by three oral and maxillofacial pathologists (JB, GB, and MM). The diagnoses included hyperkeratosis (no dysplasia), low-grade dysplasia, high-grade dysplasia, carcinoma in situ (CIS) or CIS suspicious for invasion, invasive squamous cell carcinoma, or none of the above. The final diagnosis was established based on concordance among 2/3 or 3/3 pathologists. Cases diagnosed with CIS suspicious for invasion and invasive SCC were grouped as squamous cell carcinoma.

### Tissue microarray construction and Nanostring GeoMx analysis

Hematoxylin and eosin (H&E)-stained slides were used to identify areas containing epithelium and connective tissue. A tissue microarray with 126 samples was constructed using 0.6-mm cores of FFPE tissue blocks. FFPE sections (4 µm) were cut and placed on Superfrost Plus Microscope Slides (Fischer Scientific, 12-550-15). Slide preparation was performed according to the manufacturer’s protocol (Nanostring GeoMx) with the following digestion and heat-induced epitope retrieval (HIER) conditions: 0.1 µg/ml Proteinase K for 15 min and ER for 20 min. ROI selection (PanCK+/PanCK-) was performed in the GeoMx Digital Spatial Profiler (DSP) based on morphology and immunofluorescence staining with PanCK (Novus Biologicals, clone AE1/AE3, NBP2-33200AF488), SMA (Abcam, clone 1A4, ab202368), CD45 (CST, clone D9M8I, 13917BF), and SYTO 83 DNA staining (ThermoFisher, S11364). The TMA slide was hybridized with NanoString Human Whole Transcriptome Atlas protein-coding probes, which were photocleaved and collected in 96 well-plates. Finally, a library was constructed, and sequencing was performed using an Illumina NovaSeq 6000 with S4 v1.5 100 cycle flow cell reagents and paired-end reads.

### Readout and quality assessment

Quality control was performed for each ROI. Segments were excluded according to the following criteria: <100 raw sequencing reads, <80% aligned, <80% trimmed or <80% stitched sequencing reads, <50% sequencing saturation, and <20 nuclei counts. The segment limit of quantification was calculated using the following formula: LOQi = geomean (NegProbei) * geoSD(NegProbei)2, and segments with a gene detection rate <10% were excluded. Additionally, genes detected in less than 10% of the segments were excluded from the analysis. Quartile 3 normalization was used for the downstream analyses. A total of 153 out of 160 ROIs passed quality control, while seven segments were excluded due to insufficient sequencing quality and gene detection rate. A total of 12,279 genes were detected in at least 10% of the segments and were subsequently analyzed.

### Differential gene expression

Differential gene expression (DGE) analysis was performed for eight PanCK+ and PanCK-comparisons: progressors (bx 1) versus non-progressors (bx 1), progressors (bx 1) versus progressors (bx 2), non-progressors (bx 1) versus non-progressors (bx 2), and inflamed non-dysplastic versus non-inflamed non-dysplastic specimens. DGE was evaluated on a per-gene basis by modeling normalized gene expression using a linear model with GeoMxTools v3.0.1. A Benjamini-Hochberg false discovery rate (FDR) correction was applied to the p-values. Differentially expressed genes between groups were considered significant at nominal p<0.05.

### Pathway analyses

Gene sets were defined using the KEGG BRITE database. The R GSVA package (Bioconductor version 3.15) was used to score each segment (Castelo et al., 2022). Scores were calculated for any gene set with at least five genes and fewer than 500 genes. Significant gene enrichment was established at p<0.05 and a z-score fold change of 0.5. Pathway enrichment was considered significant at p<0.05 (nominal). A second pathway enrichment analysis was performed using PathDIP 5, according to previously published protocols (Pastrello et al., 2023). The input for each comparison consisted of differentially expressed (DE) genes with nominal p<0.05 and log2fold change |0.7|.

### Spatial deconvolution

Spatial deconvolution analysis was performed using the R SpatialDecon package (Bioconductor version 3.15) (Griswold & Danaher, 2022) using a pre-specified cell profile matrix generated from scRNA-seq data to deconvolute GeoMx data, as previously described by Danaher et al. (Danaher et al., 2022).

### Cell lines

UM-SCC1 (University of Michigan Squamous Cell Carcinoma) cells were obtained from ATCC and maintained in Dulbecco’s Modified Eagle Medium (DMEM) supplemented with 10% fetal bovine serum (FBS) and 1% penicillin-streptomycin. Cells were cultured in a humidified incubator at 37°C with 5% CO□. MOC2 is a Mouse Oral Cancer (MOC) cell line derived from carcinogen-induced oral tumors in C57BL/6 mice (EWL002-FP, Kerafast) and maintained in IMDM with supplemented EGF, insulin, and hydrocortisone as described by the manufacturer’s protocol.

### siRNA- based validation of genes of interest

A custom siGENOME plate targeting all significant differentially expressed genes (DEG) was purchased from Dharmacon, including: ACTN1, AIM2, BID, BMP2, BSPRY, C10orf99, C1R, C1S, CALML3, CAPNS2, CDKN2A, CHI3L1, CLCF1, COL1A1, COL3A1, COMP, CRIP1, CTHRC1, CXCL1, DBI, DHCR24, DKK, DPYD, DSC2, DSG1, DSP, DTX3L, EMP1, FRZB, FST, HR, IFI44L, KLF4, KRT14, KRT6A, LIPM, MDK, MMP1, MMP10, MMP13, MYADM, MYCL, MYO1B, NOTCH3, PDPN, PIM2, PKP1, PLAU, POSTN, RTP4, S100A10, S100A14, SAA2, SLFN5, SOCS3, SOD2, STMN3, STOM, THBS1, TIMP1, TM4SF1, TNFRSF12A, TPPP3, TPSAB1, TRIM29, TRPM4,TXN, UBA7, ZFAT, and ZNF750. This custom plate was reconstituted in 32 μL of 1× resuspension buffer per well, yielding a final siRNA concentration of 3.125 μM. Plates were gently mixed to avoid bubble formation, incubated at room temperature on an orbital shaker for 70-90 minutes, briefly centrifuged to collect the contents at the bottom, sealed, and stored at -20 °C until use.

Prior to transfection, daughter plates were prepared by combining 0.1 μL DharmaFECT reagent (DFR) with 22.9 μL serum-free, antibiotic-free medium (SFAF) for a total volume of 23 μL per well. Subsequently, 2 μL (6.25 pmol) of siRNA was aliquoted from the master plate into each well. After incubating the mixture at room temperature for 30 min to allow complex formation, 100 μL of an antibiotic-free cell suspension was added to each well, bringing the total volume to 125 μL. Plates were then incubated at 37 °C in 5% CO for 48 h prior to downstream assays.

### Wound Healing Assay

UM-SCC1 cells were cultured to confluence in a 24-well plate format. The cells were cultured until they reached confluence. A sterile 200 μL pipette tip was used to create scratches in the cell monolayer. After washing, the cells were replenished with fresh medium containing 10% FBS, and phase-contrast images were captured as the starting point of the experiment (0 h). Where indicated, TWEAK treatment was reapplied after scratching. The cells were incubated in a 5% CO2 incubator for an additional 12 h, after which endpoint images were acquired. Wound closure was quantified by comparing the wound area at 0 and 12 h using a custom macro in FIJI (ImageJ). At each time point, the wound area was manually traced and measured to assess cell migration, and the data were analyzed using an independent two-sample t-test. Phase-contrast images were captured immediately after scratch formation (0 h) to document the initial wound area. Where indicated, UMSCC1 cells were treated with siRNA or either TWEAK (50 ng/mL) or vehicle control for 3 h.

### Invadopodium assay

Invadopodia assays and matrix degradation were performed as previously described (Magalhaes et al., 2011). Briefly, 50,000 siRNA-treated UMSCC1 cells were seeded onto 10 mm Mattek dishes coated with Alexa Fluor-488 gelatin matrix and incubated for 24 h with siRNA treatment. For the UM-SCC1 cell line, TWEAK was applied at concentrations of 50 and 100 ng/mL. For the MOC2 cell line, the TWEAK concentration was 50 ng/mL. Following incubation, the cells were fixed in 3.7% paraformaldehyde (PFA) for 20 min and subjected to immunocytochemistry for cortactin and DAPI, in accordance with previously established protocols (27). Degradation spot quantification was accomplished through the application of spot colocalization analysis (ComDet) using the ImageJ software.

### Proliferation assay

The impact of siRNA on the proliferation of corresponding genes was assessed using the alamarBlue assay (Invitrogen, catalog #DAL1025) following the manufacturer’s instructions. Briefly, UMSCC1 cells were seeded in 96-well plates at a density of 5000 cells per well, with multiple replicates. Cell density was measured on days 1, 2, and 5. AlamarBlue solution was added to each well, and the plates were incubated for 1 hour. Absorbance was then measured at 595 nm using a plate reader. he relative reduction of alamarBlue was calculated in accordance with the manufacturer’s instructions and subsequently scaled to account for technical variation between plates. An independent two-sample t-test was performed to identify significant differences in proliferation between the knockdown and control samples.

#### Statistical analysis

Statistical analysis of two variables was conducted using Student’s t-test or ANOVA with Tukey’s post hoc test for multiple groups. Using the Kaplan–Meier method, OS and DSS curves were estimated and compared using the log-rank test. Analyses were performed using GraphPad Prism and R 4.2.1 (Institute for Statistics and Mathematics of Wirtschaftsuniversität (WU), Wien, Austria: The R Project for Statistical Computing). https://www.rproject.org/). All p-values < 0.05 (two-sided) were considered statistically significant.

**Figure 1:**
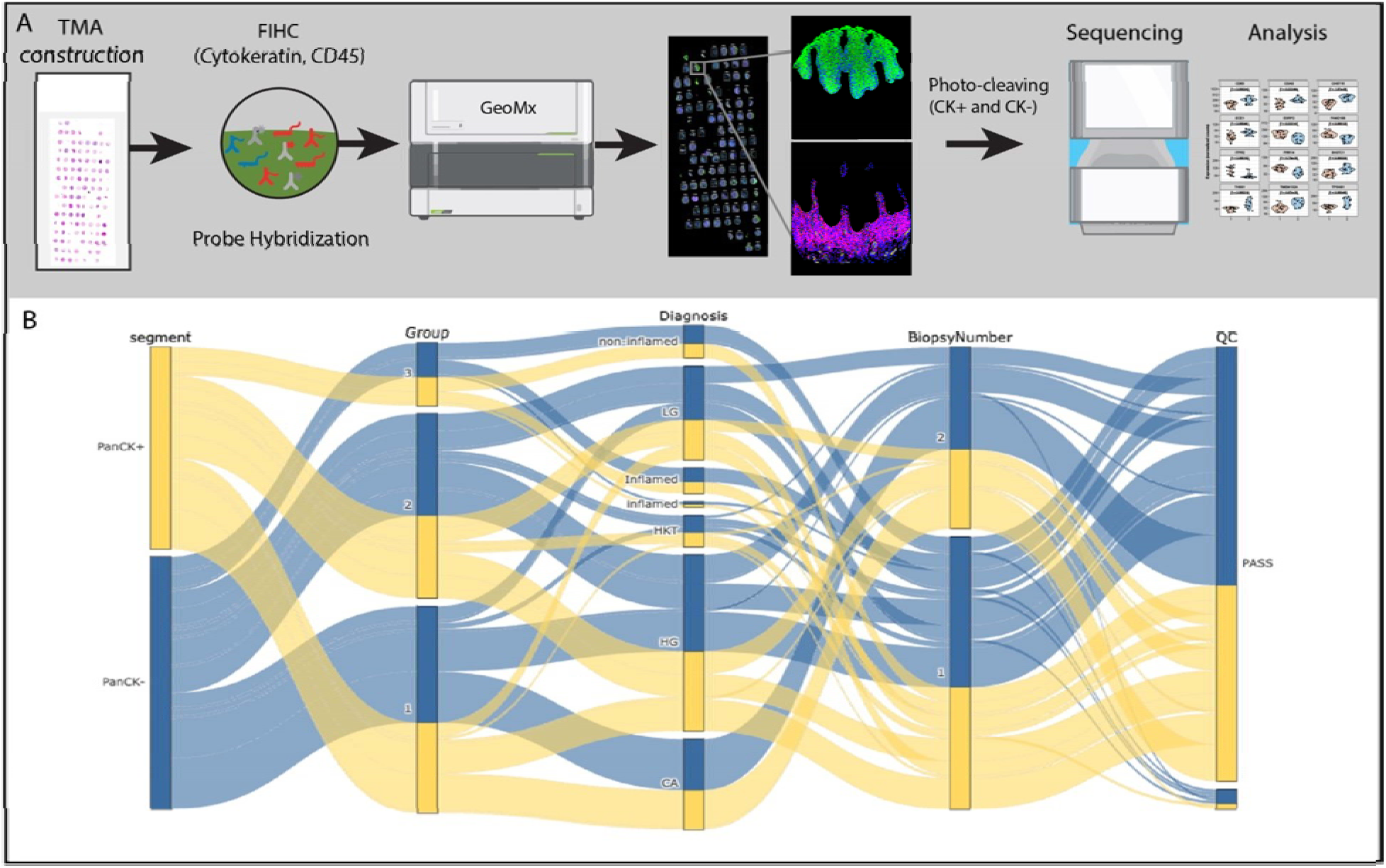
(A) Summary of methods. (B) Sankey diagram demonstrating experimental design by flow of samples through biological annotations.

## Results

Based on the selection criteria and pathology review, the following cases were included for tissue microarray construction: 24 progressing pairs, 26 non-progressing pairs, and 16 non-dysplastic specimens. A total of 126 cores were included in the TMA, of which 99 were retained after sectioning. Of the 160 PanCK+ and PanCK- ROI generated, 153 passed quality control and were included in further analyses. The clinicopathological features of the analyzed cases are summarized in Table 1. The mean time to MT was 4.47 years (range: 0.7-12.4 years) in the progressing group.

### DEG Comparison 1: Progressors (bx 1) versus non-progressors (bx 1)

Unsupervised hierarchical analysis of PanCK+ ROI (17 progressors and 15 progressors) showed the separation of samples within two clusters, with one cluster composed exclusively of progressing specimens. Overall, 65 genes were differentially expressed, with 21 upregulated and 44 downregulated genes in progressors using a dual threshold of p<0.05 and log2fold change > |0.5|. The most upregulated gene in progressing OED was STOM, whereas CDKN2A and KIF26A were the most downregulated genes ( Figure 2). Analysis of the PanCK- ROI (21 progressors and 17 non-progressors) showed sample clustering into three clusters, with one cluster composed almost exclusively of progressing specimens. Overall, 34 genes were differentially expressed, with 13 upregulated and 21 downregulated in progressors using a dual threshold of p<0.05 and log2fold change > |0.5| **(Figure 2).** In progressive OED, the topmost upregulated genes were IGHA1, FRZB, and ZFAT, whereas EMP1, S100A10, KLF4, and EPS8L2 were the topmost downregulated genes. See the Supplementary Data for a complete list of results for each comparison.

**Figure 2.**
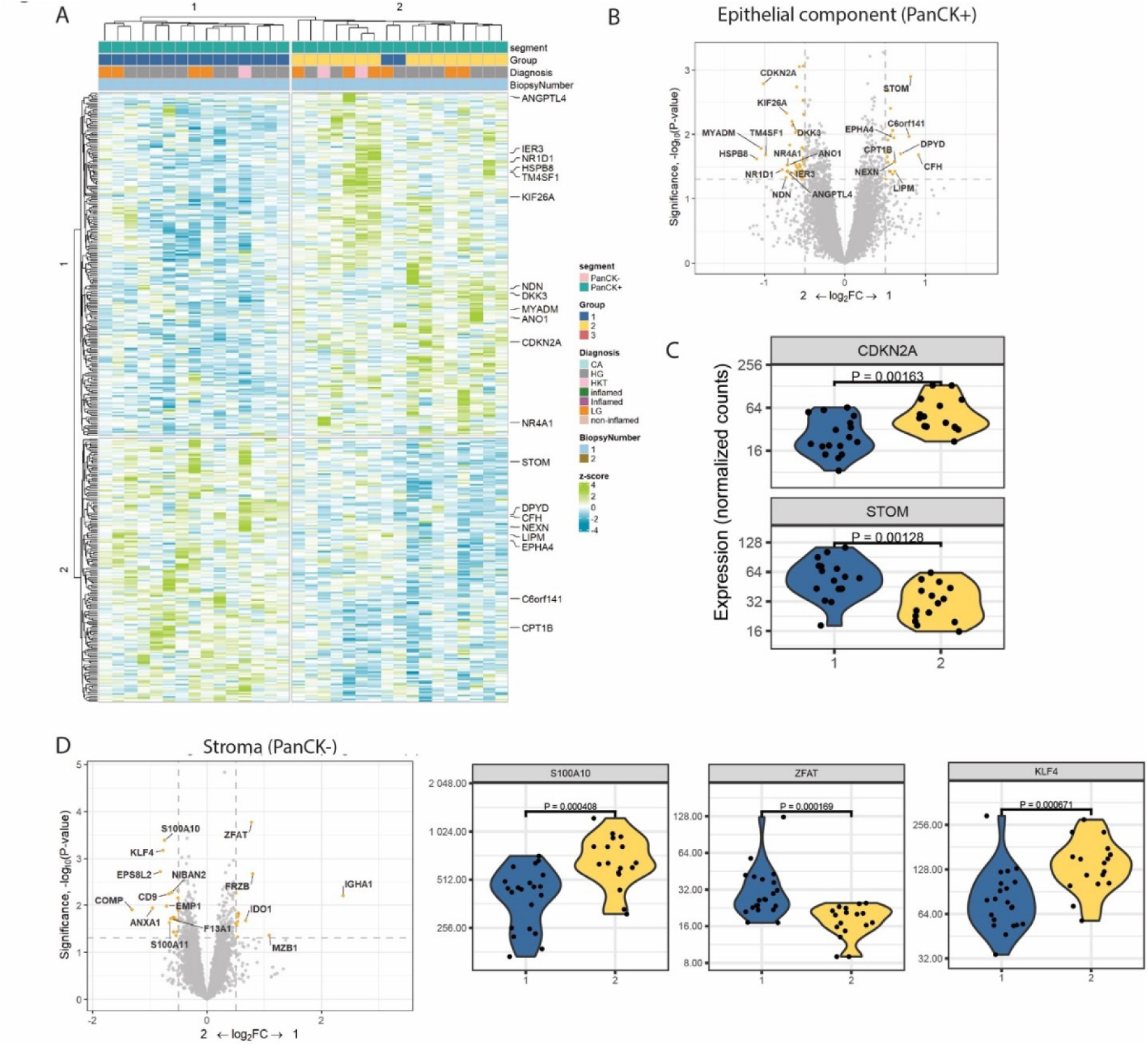
Differential gene expression in progressors (biopsy 1) and non-progressors (biopsy 1) within PanCK-positive (A-C) and PanCK-negative (D) regions of interest (ROI). (A) Heatmap depicting unsupervised clustering of samples into two groups based on differentially expressed (DE) genes. (B) Volcano plot highlighting significantly DE genes with yellow dots. (C) Violin plots illustrating the most prominently downregulated gene (CDKN2A) and upregulated gene (STOM) in progressing oral epithelial dysplasia (OED). (D) Volcano plot displaying significantly DE genes with yellow dots, alongside violin plots showing downregulated genes (S100A10 and KL4) and upregulated gene (ZFAT) in progressing OED.

### DEG Comparison 2: Progressors (bx 1) versus progressors (bx 2)

Unsupervised hierarchical analysis of PanCK+ ROI (18 precursor lesions and 13 OSCC) showed clustering into two clusters, with one cluster composed exclusively of OSCC specimens. Overall, 115 genes were differentially expressed in OSCC, with 63 upregulated and 52 downregulated, using a dual-threshold. with p<0.05 and log2fold change > |0.7**| (Figure 3).** In OSCC, the most upregulated genes were MMP1, MMP13, TPSAB1, TINAGL1, SAA2, THBS1, PRS22, SOCS3, COL1A1, and MDK, whereas TPPP3, CALML3, CAPNS2, and DSP were downregulated. DE analysis of the stroma (21 precursor lesions and 18 OSCC) revealed two clusters, one of which was predominantly composed of high-grade OED and OSCC. Overall, 31 genes were differentially expressed in OSCC, with 25 upregulated and 6 downregulated, using a dual threshold of p<0.05 and log2fold change > |0.7| **(Figure 3)**. In OSCC, the most upregulated genes were IGHM, MMP1, POSTN, PIM2, CXCL1, MT2A, TIMP1, CTHRC1, and NNMT, whereas ANK2 was downregulated.

**Figure 3.**
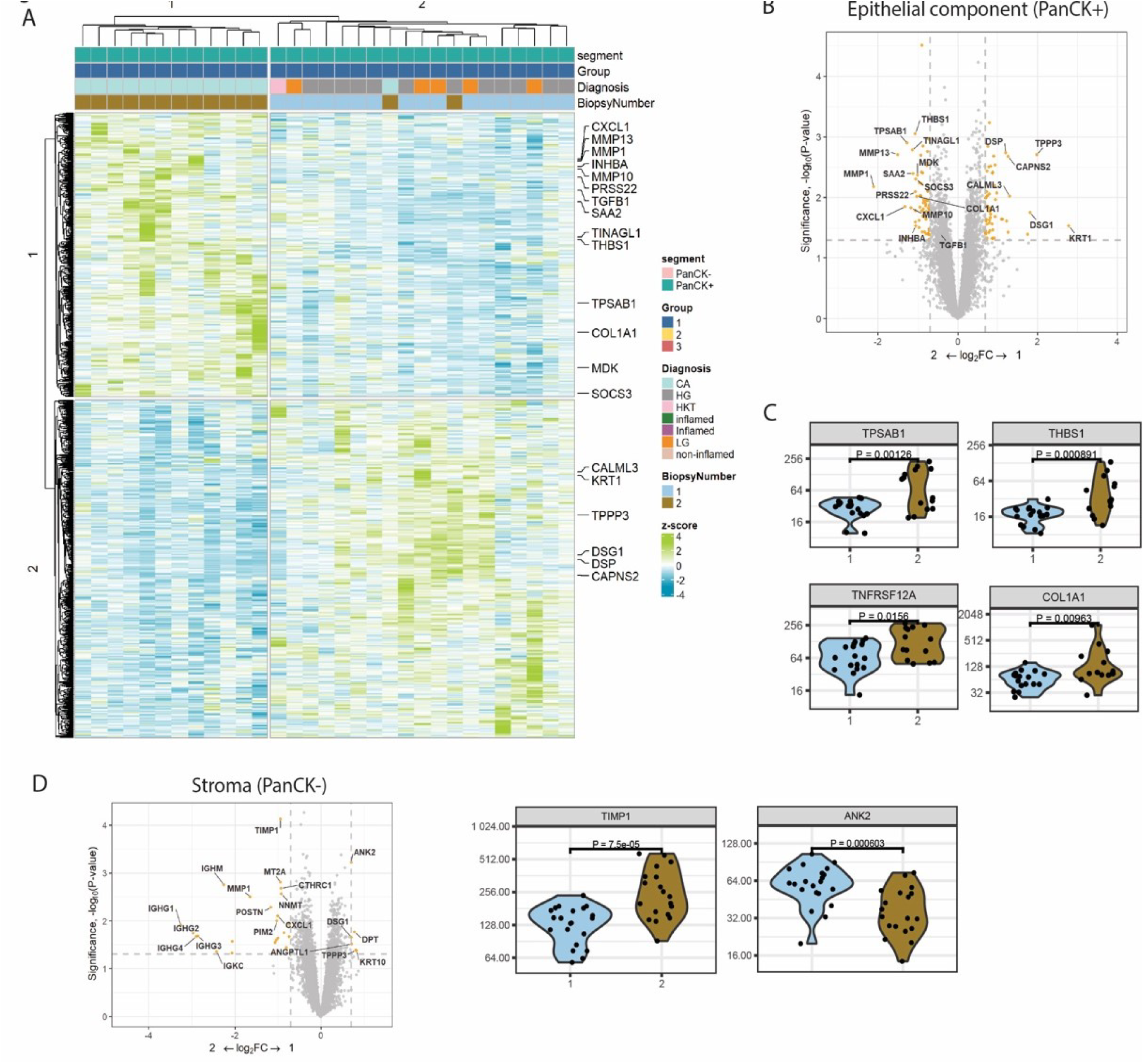
Differential gene expression in progressors (biopsy 1) and progressors (biopsy 2) (A) Heatmap depicting unsupervised clustering of samples into two groups based on differentially expressed genes. (B)Volcano plot highlighting differentially expressed genes with yellow dots. (C) Violin plots illustrating upregulated genes in progressing oral epithelial dysplasia. (D) Volcano plot indicating significantly differentially expressed genes with yellow dots, along with representative violin plots showing downregulated (ANK2) and upregulated (TIMP1) genes in oral squamous cell carcinoma (OSCC).

### Comparison 3: Non-progressors (bx 1) versus non-progressors (bx 2)

Unsupervised hierarchical analysis of the PanCK+ ROI (15 biopsy 1 and 13 biopsy 2) revealed two clusters of cells. Overall, 39 genes were differentially expressed, with 14 upregulated and 25 downregulated in biopsy 2, using a dual threshold of p < 0.05 and log2 fold change > |0.5|. Stromal analysis (PanCK-) (17 biopsy 1 and 16 biopsy 2) also showed clustering into two clusters, one of which was composed exclusively of biopsy 2 samples. Overall, 23 genes were differentially expressed, with 9 upregulated and 14 downregulated in biopsy 2, using a dual threshold of p < 0.05 and log2 fold change > |0.5|.

### Comparison 4: Inflamed non-dysplastic versus non-inflamed non-dysplastic samples

In the epithelial component (PanCK+), 77 genes were differentially expressed in inflamed samples, with 48 upregulated and 29 downregulated, using a dual threshold of p < 0.05 and log2 fold change > 0.7. A total of 799 genes were differentially expressed in the stroma, with 771 upregulated and 28 downregulated in inflamed samples, using a dual threshold of p < 0.05 and log2 fold change > 1.0.

### Pathway Analysis

Using the PathDIP pathway analysis database, we identified the most significantly enriched pathways in the progressing samples as signal transduction, cancer, immune system, and development and regeneration **(Figure 4).** Supplementary tables are provided for each comparison.

**Figure 4.**
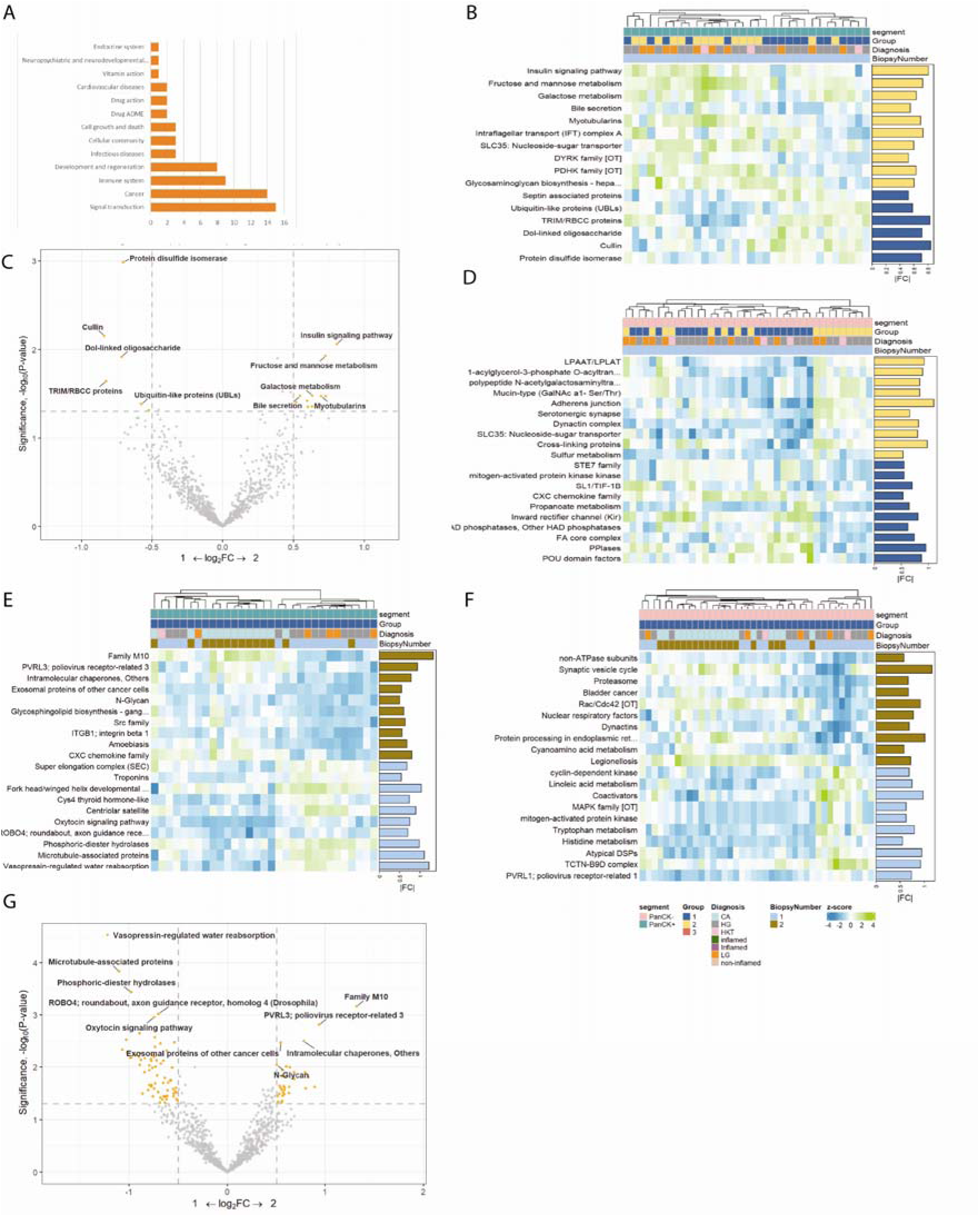
Pathways analyses. (A) Bar graph showing the most enriched pathways in Progressors, indicating the number of enriched pathways within each category. (B) Heatmap illustrating clustering of PanCK-positive (PanCK+) regions of interest (ROI) in progressing and non-progressing oral epithelial dysplasia (OED). (C) Volcano plot displaying differentially expressed (DE) pathways, with yellow dots representing significant pathways in PanCK+ ROI of progressors and non-progressors. (D) Heatmap showing clustering of PanCK-negative (PanCK-) ROI in progressing and non-progressing OED. (E) Heatmap illustrating clustering of PanCK+ ROI in precursor lesions and subsequent oral squamous cell carcinoma (OSCC) samples. (F) Heatmap depicting clustering of PanCK- ROI in precursor lesions and subsequent OSCC samples. (G) Volcano plot with yellow dots indicating DE pathways in PanCK+ ROI of precursor lesions and subsequent OSCC.

#### Comparison 1: Progressors (bx 1) versus non-progressors (bx 1)

Within the epithelial component, gene set variation analysis (GSVA) revealed eight pathways significantly enriched in progressors, whereas 13 were downregulated. The pathways enriched for upregulated genes included protein disulfide isomerase, cullin, TRIM/RBCC, and ubiquitin-like proteins (Figure 4). The pathways enriched for downregulated genes included the insulin signaling pathway, fructose and mannose metabolism, and galactose metabolism. Similarly, in the stromal component of the progressors, 12 pathways were upregulated and 28 were downregulated.

#### Comparison 2: Progressors (bx 1) versus progressors (bx 2)

GSVA revealed 46 upregulated pathways and 77 downregulated pathways enriched within the epithelial component of OSCC (Figure 4). The matrix metallopeptidase family (M10), which plays a role in epithelial-to-mesenchymal transition, was significantly represented in the results. The stromal component exhibited 49 upregulated and 30 downregulated pathways.

#### Comparison 3: Non-progressors (bx 1) versus non-progressors (bx 2)

GSVA revealed five upregulated and 12 downregulated pathways in the epithelial component of biopsy 2. The stroma showed 14 upregulated and 15 downregulated pathways, respectively.

### Spatial deconvolution

Unsupervised hierarchical clustering revealed four clusters across all ROI (Figure 5A). One cluster mainly consisted of epithelial components and showed a reduced number of inflammatory cells. The remaining three clusters mainly contained PanCK-ROI. PanCK- segments exhibited higher levels of memory CD8 T cells, macrophages, regulatory T cells, endothelial cells, and fibroblasts, among other cell types. In contrast, PanCK+ segments had lower cell diversity than PanCK- segments. Plasma cells (p=0.03) were increased in the epithelium of progressing (bx 1) samples compared to non-progressing (bx 1) samples. Fibroblasts (p=0.0057) and neutrophils (p=0.0078) were more abundant in OSCC than in the matching precursor lesions (Figure 5B). Additionally, fibroblasts (p=0.013), neutrophils (p=0.005), monocytes (p=0.029), and mast cells (p=0.045) were elevated in the PanCK- ROI of OSCC compared to their corresponding precursor lesions (Figure 5B). The analysis of the proportion estimates highlighted the substantial presence of fibroblasts in the analyzed samples (Figure 5C).

**Figure 5.**
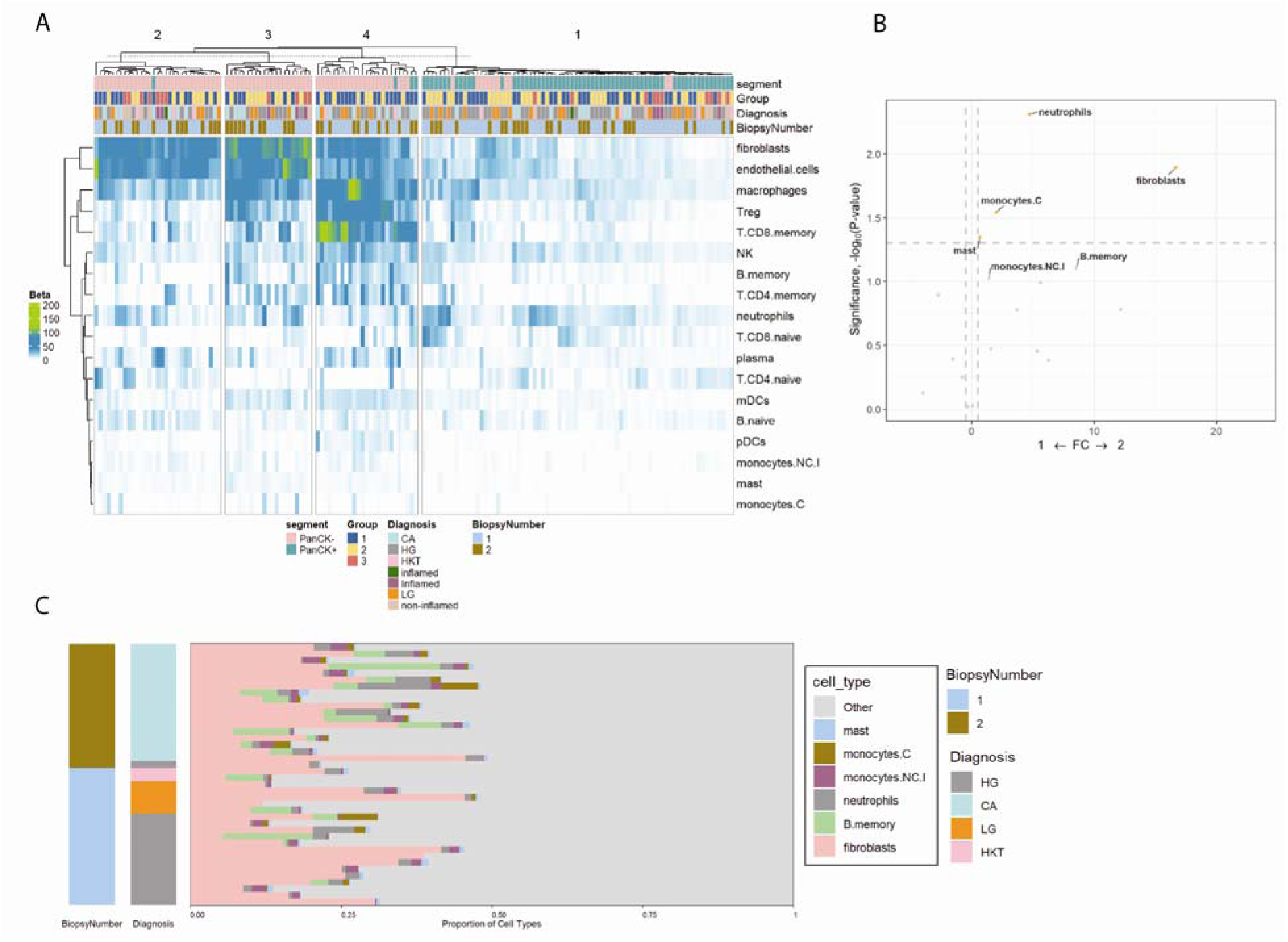
Spatial deconvolution analysis performed in this study. (A) heatmap generated via unsupervised clustering encompassing all Regions of Interest (ROIs) analyzed. Panel (B) illustrates a volcano plot identifying cell populations within the microenvironment that exhibit significant alterations when comparing progressing dysplasia to Oral Squamous Cell Carcinoma (OSCC). Specifically, the yellow dots on the plot denote cell populations with a significant increase in the OSCC microenvironment. Panel (C) presents a stacked bar plot that visually contrasts the proportions of various cell types in the microenvironment of OSCC relative to the precursor lesion, Oral Epithelial Dysplasia (OED).

### Cell line validation

Using the top differentially expressed genes (|log2FC| > 0.7, p < 0.05) across three comparisons—progressor (bx1) versus progressor (bx2), non-progressor (bx1) versus non-progressor (bx2), and progressor (bx1) versus non-progressor (bx1)—a preliminary list was generated. From this list, genes of interest, particularly those involved in cell survival, were further refined. Ultimately, 105 genes were selected for downstream validation and analysis (Figure 6A).

**Figure 6.**
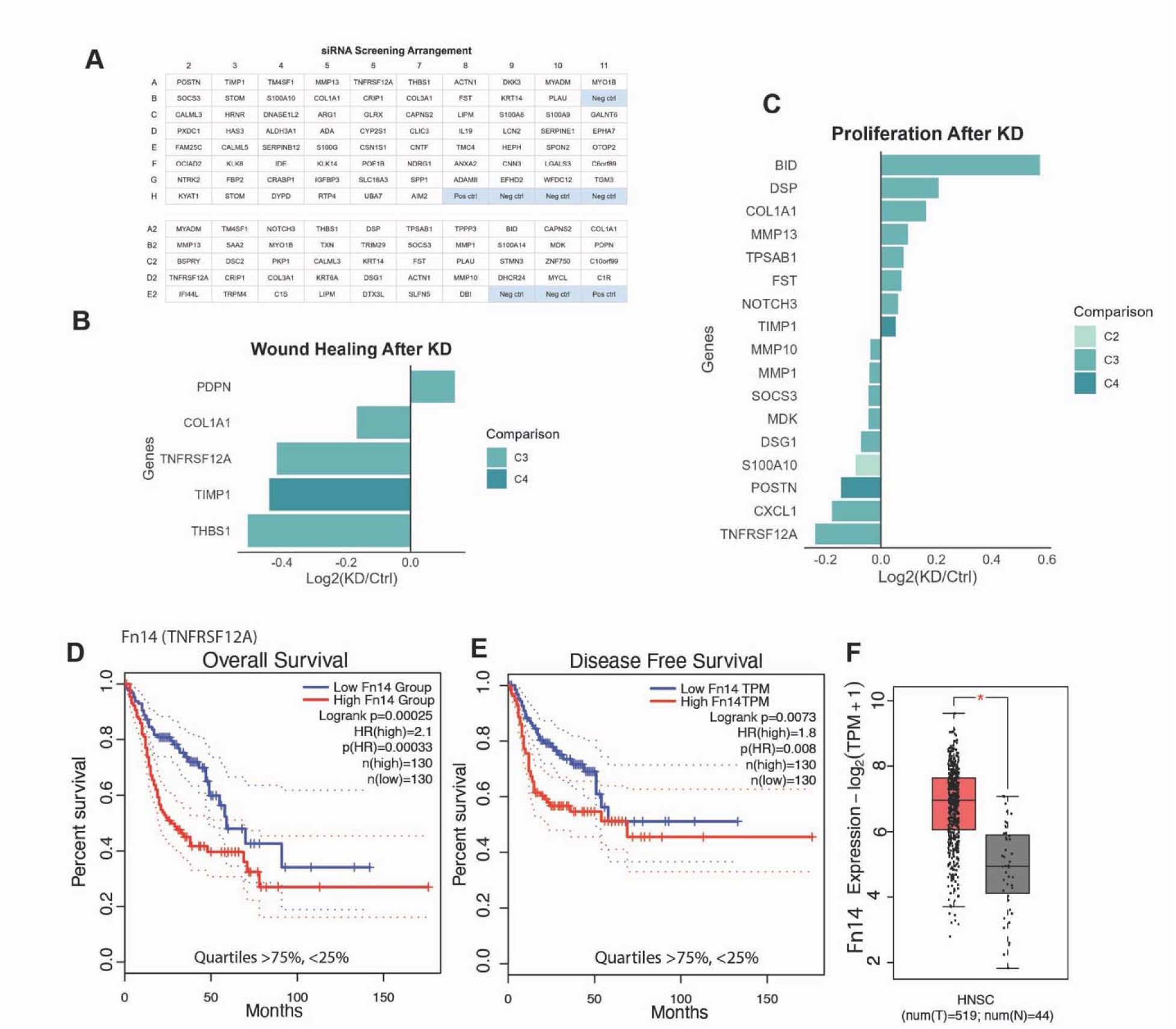
Selection of candidate genes for functional validation through proliferation, migration assays, and GEPIA analysis. Genes identified from initial transcriptomic analyses (see Figure 2 and Figure 3) (A) were selected for further analysis using siRNA-based gene silencing, followed by wound healing (B) and cell proliferation (C) assays to assess their roles in regulating cell growth and motility. The genes associated with specific comparisons are listed Comparison 2 (C2): CK- Progressors (bx 1) versus non-progressors (bx 1); comparison 3 (C3): CK+ progressors (bx 1) versus progressors (bx 2) and comparison 4 (C4): CK- progressors (bx1) versus progressors (bx2). The remaining comparisons had no significant genes on these assays. N=2, P<0.05 on both replicates. Overall survival (D), disease-free survival (E), and expression (F) of TNFRSF12A using The Cancer Genome Atlas (TCGA) dataset via GEPIA (Gene Expression Profiling Interactive Analysis).

An siRNA screen was developed to investigate the role of these genes in cell migration and proliferation. Knockdown of the following genes significantly affected cell migration, as demonstrated in the wound healing assay: PDPN, TNFRSF12A, COL1A1, THBS1, and TIMP1 (Figure 6B). All knockdowns reduced migration, except for PDPN.

Knockdown of S100A10, TNFRSF12A, MDK, SOCS3, MMP10, CXCL1, MMP1, DSG1, and POSTN reduced proliferation. In contrast, knockdown of BID, FST, COL1A1, TPSAB1, MMP13, DSP, NOTCH3, and TIMP1 increased proliferation (Figure 6C)

Expression levels and their correlations with overall survival were analyzed using The Cancer Genome Atlas (TCGA) dataset through Gene Expression Profiling Interactive Analysis (GEPIA). The analysis demonstrated that overexpression of TIMP1 (see Supplemental Figure 1), THBS1, and particularly TNFRSF12A was associated with reduced overall survival (Figure 6D) and disease-free survival (Figure 6E). Additionally, these genes were significantly overexpressed in tumor tissues from head and neck cancer patients with HNSCC (Figure 6F).

#### TWEAK, a cytokine that activates FN14 signaling, promotes migration, matrix degradation

Activation of Fn14 by TWEAK enhances cell migration. As shown in Figure 7A, Treatment with TWEAK (50 ng/mL) significantly increased cell migration, with wound healing increasing from 55.8% in the control group to 98.84% in the treated group (42%, p = 0.00047). As shown in Figure 7C, Fn14 knockdown significantly impaired the migratory capacity of the UMSCC1 cells. Similarly, the invadopodia formation assay revealed a notable reduction in matrix degradation following Fn14 silencing (Figure 7D). Figure 7E-F illustrate that TWEAK stimulation at 100 ng/mL significantly enhanced matrix degradation, with an average degraded area of 276.2 µm², representing a 12-fold increase compared to the control (21.65 µm², p = 0.0006). A significant difference was also observed between the 50 ng/mL and 100 ng/mL TWEAK-treated groups (p = 0.0228). Although matrix degradation increased to 106.0 µm² at 50 ng/mL, the difference was not statistically significant compared to that in the control group (p = 0.3571). TWEAK treatment also promoted cell invasion, as demonstrated by the invasion assay (Figure 7G). Data were analyzed using one-way ANOVA, followed by Tukey’s multiple comparisons test.

**Figure 7.**
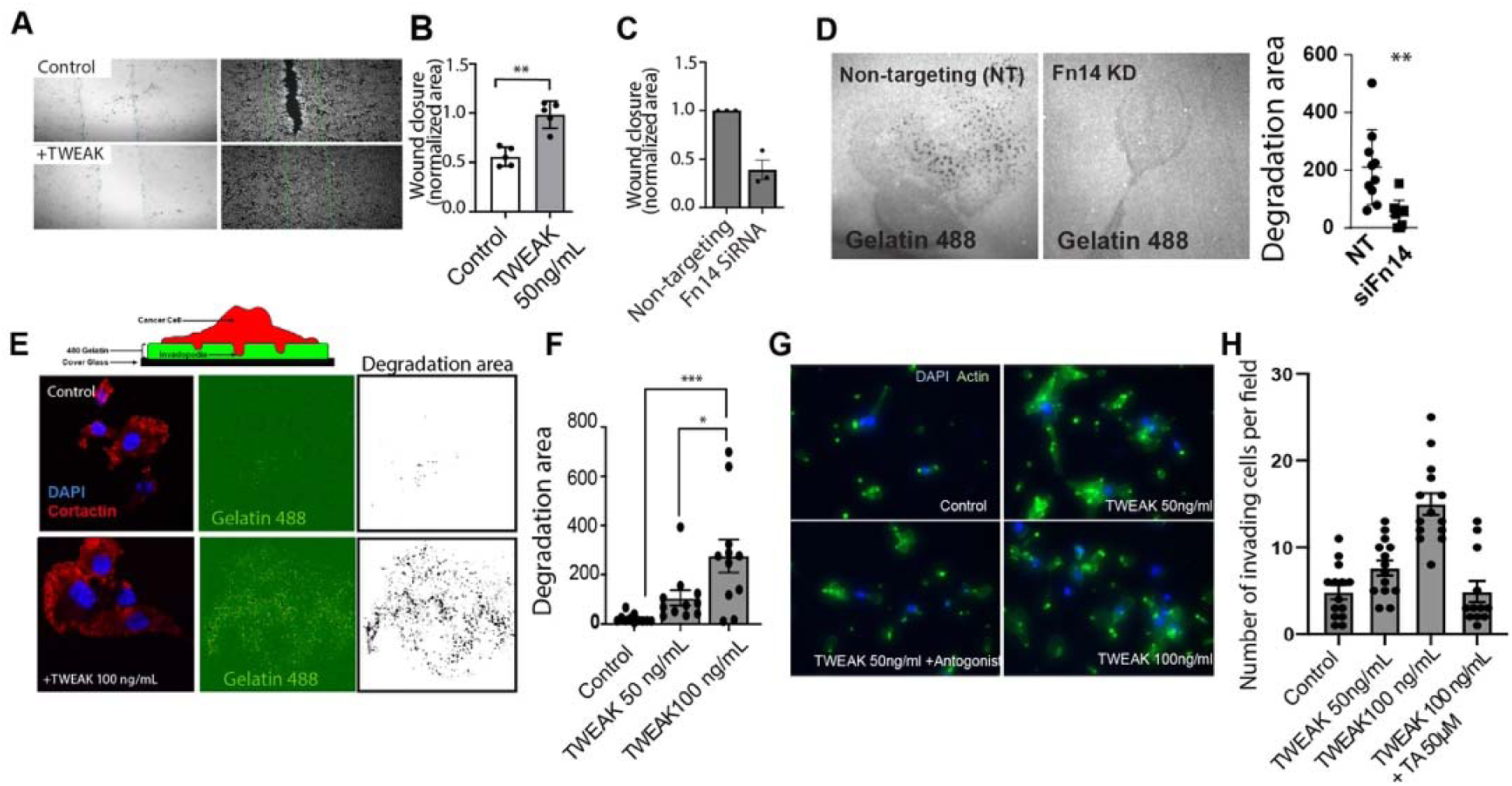
TWEAK induces migration and invasion of UMSCC1 cells. (A) Representative images of a wound healing assay showing the migration of UMSCC1 cells with or without TWEAK stimulation. (B) Quantification of wound closure 12 hours after TWEAK stimulation at 50 ng/mL (n=3, p<0.5). (C) Quantification of UMSCC1 cell wound closure following Fn14 siRNA Silencing effects compared to a non-target siRNA control (n=3, p<0.05). (D) Representative images of the invadopodia formation assay evaluating invadopodia-dependent matrix degradation, indicated by loss of fluorescence, following stimulation with TWEAK at 50 ng/mL and 100 ng/mL. UMSCC1 cells were stained with Cortactin Alexa555 and DAPI, with the gelatin matrix containing Gelatin Oregon Green 488 for visualization. Quantification of matrix degradation after TWEAK stimulation of UMSCC1 cells (E) and after siRNA-mediated silencing of Fn14 in UMSCC1 cells (n=3, p<0.05) (F). (G) Representative images of the transwell invasion assay, showing the underside of the membrane stained with DAPI and Phalloidin 488. Quantitative analysis is presented in the corresponding graph (H). Data represent mean ± SEM from three independent experiments. A p-value < 0.05 was considered statistically significant, with TWEAK at 100 ng/mL.

#### Fn14 is overexpressed in progressing oral dysplasia

To evaluate Fn14 expression in oral lesions with distinct clinical outcomes, we constructed an additional tissue microarray (TMA) comprising samples from four patients with progressing dysplasia (eight samples, collected before and after cancer development) and four patients with non-progressing dysplasia (eight samples, each with two sequential biopsies showing no progression). Fn14 expression was assessed across these groups by comparing non-progressing dysplasia, progressing dysplasia, and oral squamous cell carcinoma (OSCC). Our results demonstrated an increase in Fn14 expression in the carcinoma group compared with that in the dysplasia group (Figure 8B and 8C).

**Figure 8:**
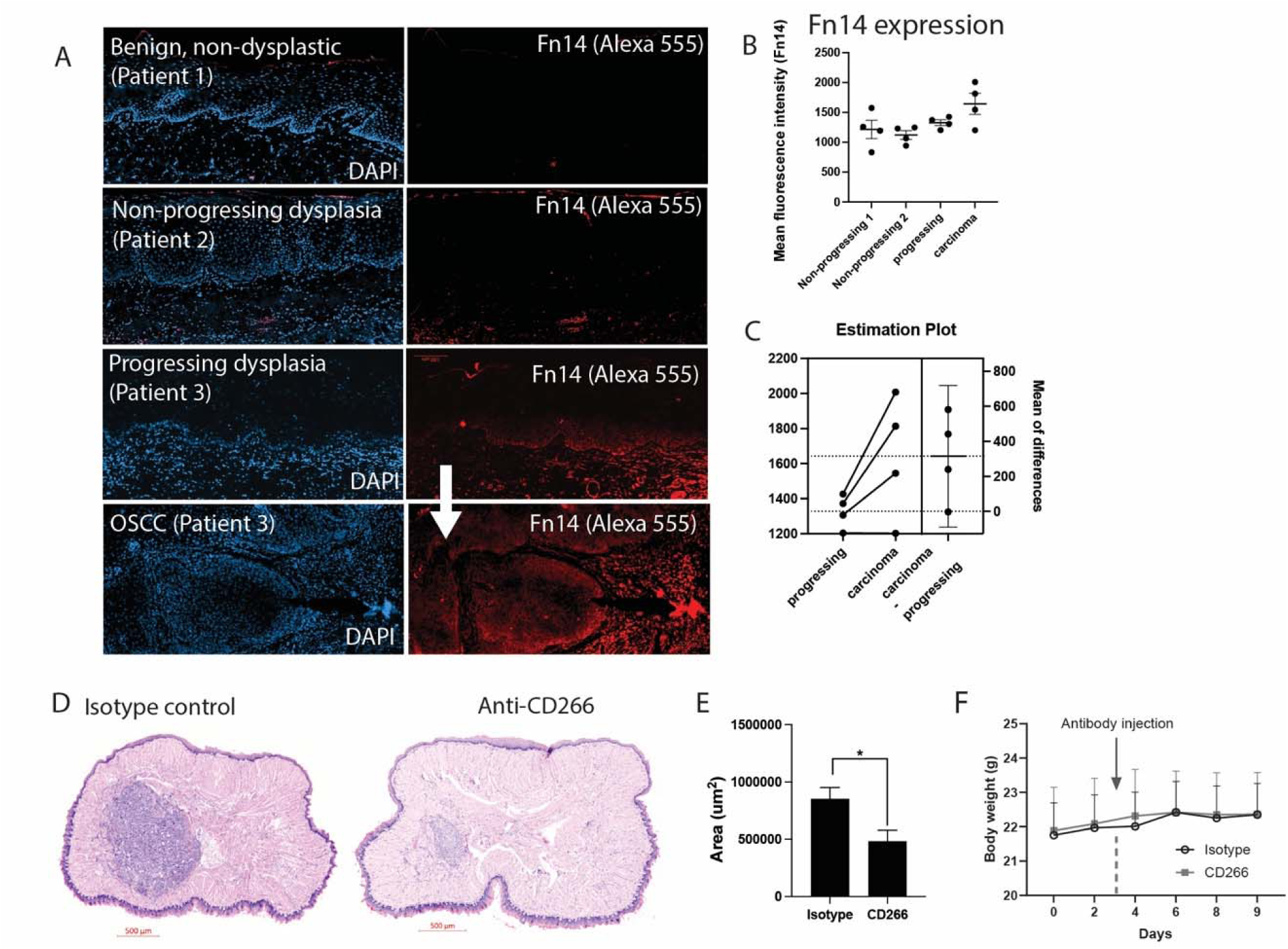
(A) Representative fluorescent immunohistochemistry (IHC) photomicrographs illustrating Fn14 (Alexa 555) expression in paired samples of progressing dysplasia and squamous cell carcinoma (SCC) from patients 1 and 2, as well as non-progressing cases from patient 3. A total of 16 specimens, derived from 8 patients (2 biopsies per patient), were analyzed, and the Fn14 expression data are summarized in panel B (n=8, p<0.05). Additionally, representative Hematoxylin and Eosin (H&E)-stained photomicrographs depict the tongues of mice injected with MOC2 cells. Panel C compares animals administered with an isotype control (50 μg/mouse, weekly injections) to those receiving anti-CD266 neutralization (50 μg/mouse, weekly injections). Panel E displays body weight of 4NQO-treated mice over time. (F) tumor volume quantification is presented in Panel F). (n=9 per group) Experimental data are presented as mean ± SEM. *p < 0.05.

### Fn14 neutralization reduces tumour size in an allograft oral cancer model

To assess the therapeutic potential of targeting CD266 in oral cancer, we used an MOC2 oral tongue allograft model. Administering anti-CD266 (50 μg/mouse) significantly inhibited tumor growth compared to the isotype control group (p = 0.05) (Figure 8D, 8E). Throughout the experiment, the body weight remained stable, with no significant weight loss observed, indicating that the treatment was generally well tolerated (Figure 8F). Histological examination of H&E-stained sections confirmed a reduced tumor burden in the anti-CD266–treated group compared to that in the control group. These findings were consistent across independent replicates (n = 9 mice per group).

## Discussion

OPMDs exhibit an intrinsic risk of malignant transformation (MT) into oral squamous cell carcinoma (OSCC). While clinicopathological parameters—including lesion appearance, patient social history, and histopathological grading of oral epithelial dysplasia (OED)—remain vital for guiding patient management, their predictive accuracy for identifying high-risk lesions is limited. Consequently, many patients undergo extended surveillance and require specialized care, imposing clinical and psychological burdens.

The molecular mechanisms driving the progression of OPMDs to OSCC are complex and involve dynamic alterations in gene expression, signaling pathways, and tumor microenvironment composition. Comprehensive molecular analysis holds promise for enhancing prognostication and enabling earlier, more targeted interventions. To our knowledge, this study is the first to employ spatially resolved transcriptomic profiling on formalin-fixed, paraffin-embedded (FFPE) tissues encompassing OED, OSCC, and non-dysplastic mucosa. By analyzing matched progressing and non-progressing OED specimens, we identified transcriptomic signatures and microenvironmental features linked to malignant transformation, elucidating molecular events predating OSCC development.

Results demonstrated a clear segregation of progressing versus non-progressing dysplastic epithelium into two distinct clusters. Additionally, comparison between precursor lesions and subsequent OSCC revealed nearly perfect clustering, underscoring the potential of spatial transcriptomics for OED prognostication, contingent upon validation in independent cohorts.

Differential gene expression analysis indicated upregulation of STOM and downregulation of CDKN2A and KIF26A as common markers in epithelium of progressing OED. Reduced CDKN2A expression—a tumor suppressor involved in cell cycle regulation—is a well-established event in oral precancer and OSCC (Bradley et al., 2010; Shahnavaz et al., 2001; The Cancer Genome Atlas Network, 2015; William et al., 2016, 2019; Zhang et al., 2012). Stomatin, a transmembrane ion transporter implicated in prostate tumor suppression, has not been previously studied in oral carcinogenesis (Rahman et al., 2021). Our data suggest that downregulation of CDKN2A and upregulation of STOM and KIF26A may serve as early molecular markers distinguishing progressing from non-progressing OED. Functional assays following gene knockdown demonstrated that cell migration and proliferation are modulated by these genes, consistent with existing literature. For example, BID knockdown resulted in increased proliferation, aligning with its role in apoptosis regulation (Köhler et al., 2008). Conversely, MDK knockdown decreased proliferation, supporting its tumor-promoting function (Filippou et al., 2020). Additionally, knocking down THBS1 reduced migration, consistent with its pro-migratory role in tumors including OSCC (Pal et al., 2016).

Notably, knockdown of TNFRSF12A (TWEAKR/Fn14) decreased both proliferation and migration, suggesting its oncogenic role in OSCC. TNFRSF12A activates downstream pathways such as NF-κB, and elevated expression correlates with tumor aggressiveness, invasion, metastasis, and angiogenesis (Winkles, 2008).

Pathway analysis of progressing OED epithelium revealed upregulation of ubiquitin-like proteins, cullins, TRIM/RBCC proteins, and protein disulfide isomerase—linked to carcinogenesis via mechanisms involving ubiquitination, cell cycle regulation, DNA repair, and tumor suppressor modulation (Cambiaghi et al., 2012; Hoeller et al., 2006; Lee & Zhou, 2010; Zhao et al., 2021). Prior studies have associated DNA damage repair pathways with OED progression (Farah et al., 2019), indicating potential therapeutic targets.

Spatial deconvolution analyses showed increased plasma cell infiltration within progressing OED epithelium. Although the significance remains uncertain due to limited data on intraepithelial inflammatory cells in the esophagus, stromal analysis indicated elevated fibroblasts, neutrophils, monocytes, and mast cells, implicating cancer-associated fibroblasts and neutrophils in malignant transformation (Bienkowska et al., 2021; Glogauer et al., 2015; Magalhaes et al., 2014).

Limitations include challenges in constructing tissue microarrays from small oral samples while maintaining tissue architecture. Of 126 specimens, 99 (78%) were suitable for evaluation after sectioning. Technical limitations inherent to transcriptomics, such as difficulty quantifying low-abundance transcripts in small regions (<100 µm), were observed. No genes reached significance after false discovery rate correction, likely due to sample size. Ongoing validation studies in independent cohorts aim to confirm these molecular findings and their relevance to proteomic and biological outcomes.

## Acknowledgements

We acknowledge Cecilia Cabral-Bolarinho from Mount Sinai Services and Dhaarmini Rajshankar from the Collaborative Advanced Microscopy Laboratories of Dentistry (CAMiLoD) of the Faculty of Dentistry, University of Toronto, Toronto, ON, Canada, for the service, training, and expert advice received on histology and microscopy imaging. This study was supported by the University of Toronto Oral Pathology Graduate Student Research Fund and CIHR project grant (PJT-17512) to Marco Magalhaes

## Data availability

The datasets generated and/or analysed during the current study will be available (to be uplodaded)

## Author Contributions

Vincent Lavoie – Data curation, data analysis, manuscript preparation

Will Jeong – Data analysis and manuscript preparation

James Jeon - Data curation and analysis

Aiman Ali – Data curation, analysis, and manuscript preparation

Juan F. Andrade - Data curation and manuscript review

Graziella Rigueira Molska - Data curation, analysis and manuscript review

Najmeh Esfandiari – Data analysis and manuscript review

Hui Ling Yeo - Data curation, analysis and manuscript review

Igor Jurisica - Data analysis and manuscript review

Deepika Chugh – Data curation and manuscript review

Iona Leong - Data curation and manuscript review

Justin Bubola - Data curation and manuscript review

Grace Bradley - Data curation and manuscript review

Marco Magalhaes – conceptualization, supervision, data analysis, manuscript preparation

