## Supplementary figures and images for "Spatial transcriptomic analysis of progressing oral epithelial dysplasia reveals unique differentially expressed genes and microenvironmental changes"

### Supplemental 1

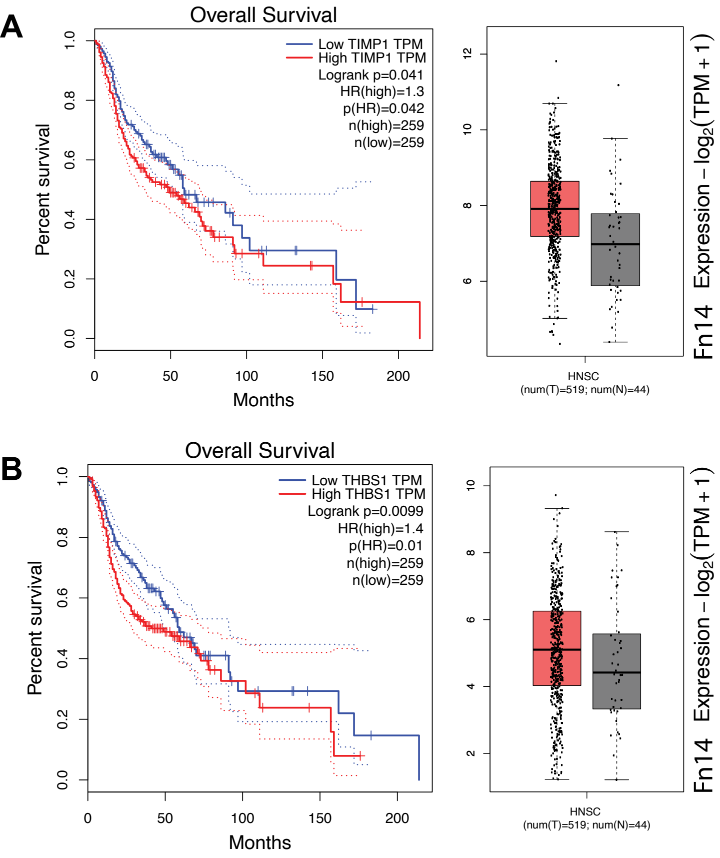
